# A dynamic nomenclature proposal for SARS-CoV-2 to assist genomic epidemiology

**DOI:** 10.1101/2020.04.17.046086

**Authors:** Andrew Rambaut, Edward C. Holmes, Verity Hill, Áine O’Toole, JT McCrone, Chris Ruis, Louis du Plessis, Oliver G. Pybus

## Abstract

The ongoing pandemic spread of a novel human coronavirus, SARS-COV-2, associated with severe pneumonia disease (COVID-19), has resulted in the generation of thousands of virus genome sequences. The rate of genome generation is unprecedented, yet there is currently no coherent nor accepted scheme for naming the expanding phylogenetic diversity of SARS-CoV-2. We present a rational and dynamic virus nomenclature that uses a phylogenetic framework to identify those lineages that contribute most to active spread. Our system is made tractable by constraining the number and depth of hierarchical lineage labels and by flagging and declassifying virus lineages that become unobserved and hence are likely inactive. By focusing on active virus lineages and those spreading to new locations this nomenclature will assist in tracking and understanding the patterns and determinants of the global spread of SARS-CoV-2.

---

There are currently more than 5000 publicly available complete or near-complete genome sequences of SARS-CoV-2 (as of 10th April 2020) and the number rises daily. This remarkable achievement has been made possible by the rapid genome sequencing and online sharing of SARS-CoV-2 genomes by public health and research teams worldwide. These genomes have the potential to provide invaluable insights into the ongoing evolution and epidemiology of the virus during the global pandemic and will likely play an important role in its eventual mitigation and control. Despite such a wealth of data, there is currently no coherent system for naming and discussing the growing number of phylogenetic lineages that comprise the population diversity of this virus, with conflicting *ad hoc* and informal systems of virus nomenclature in circulation. A nomenclature system is urgently required before scientific literature and communication become further confused.

There is no universal approach to classifying virus genetic diversity below the level of a virus species. Typically, genetic diversity is categorised into distinct ‘clades’, each of which corresponds to a monophyletic group on a phylogenetic tree. These clades may be referred to by a variety of terms, such as ‘subtypes’, ‘genotypes’, ‘groups’, depending on the taxonomic level under investigation or the established scientific literature for the virus in question. The clades usually reflect an attempt to divide pathogen phylogeny and genetic diversity into a set of groupings that are approximately equally divergent, mutually exclusive and statistically well supported. All genome sequences are therefore allocated to one clade or provisionally labelled as unclassified. Often multiple hierarchical levels of classification exist for the same pathogens, such as the terms ‘type’, ‘group’ and ‘subtype’ that are used in the field of HIV research.

Such systems are useful for discussing epidemiology and transmission when the number of taxonomic labels remains roughly constant through time; this is the case for slowly evolving pathogens (e.g. many bacteria) and for rapidly evolving viruses with low rates of lineage turnover (e.g. HIV and HCV). In contrast, some rapidly evolving viruses such as influenza A are characterised by high rates of lineage turnover, so that the genetic diversity circulating in any particular year largely emerges out of and replaces the diversity present in the preceding few years. For human seasonal influenza, this behaviour is the result of strong natural selection among competing lineages. In such circumstances a more explicitly phylogenetic classification system is used; for example, avian influenza viruses are classified into ‘subtypes’, ‘clades’ and ‘higher order clades’ according to several quantitative criteria (WHO/OIE/FAO H5N1 Evolution Working Group 2012). Such a system can provide a convenient way to refer to the emergence of new (and potentially antigenically distinct) variants and is suitable for the process of selecting the component viruses for the regularly updated influenza vaccine. A similar approach to tracking antigenic diversity may be needed to inform SARS-CoV-2 vaccine design efforts. While useful, we recognise that dynamic nomenclature systems based on genetic distance thresholds have the potential to over-accumulate cumbersome lineage names.

In an ongoing and rapidly changing epidemic, such as SARS-CoV-2, a nomenclature system can facilitate real-time epidemiology by providing commonly agreed labels to refer to viruses circulating in different parts of the world, thereby revealing the links between outbreaks that share similar virus genomes. Further, a nomenclature system is needed to describe virus lineages that vary in phenotypic or antigenic properties (although it must be stressed that at present there is no evidence of such variation among currently available SARS-CoV-2 strains).

There are a number of key challenges in this endeavour, and to be valid and broadly accepted a nomenclature needs to: (i) capture local and global patterns of virus genetic diversity in a timely and coherent manner, (ii) track emerging lineages as they move among countries and between populations within each country, (iii) be sufficiently robust and flexible enough to accommodate new virus diversity as it is generated, and (iv) be dynamic, such that it is able to incorporate both the birth and death of viral lineages through time.

A special challenge in the case of COVID-19 is that genome sequence data is being generated rapidly and at high volumes, such that by the end of the pandemic we can expect tens of thousands of SARS-CoV-2 genomes to have been sequenced. Any lineage naming system must therefore be capable of handling tens to hundreds of thousands of virus genomes sampled longitudinally and densely through time. Further, to be practical, any lineage naming system should have no more than one or two hundred active lineage labels, as any more would obfuscate rather than clarify discussion and will be difficult to conceptualise.

To fulfil these requirements we propose a workable and practical lineage nomenclature for SARS-CoV-2 that arises from a set of fundamental evolutionary and phylogenetic principles. Some of these principles are, necessarily, specific to the COVID-19 pandemic, reflecting the new normal of large-scale real-time generation of virus genome sequences. The nomenclature system is not intended to represent every evolutionary change in SARS-CoV-2, as these will number many thousand by the end of the pandemic. Instead, the focus is on genetic changes associated with important epidemiological and biological events. Fortunately, because of the early sampling and genome sequencing of COVID-19 cases in China, especially in Hubei province, it appears that the “root sequence” of SARS-CoV-2 is known. Many of the genomes from the earliest sampled cases are genetically identical and hence also likely identical to the most recent common ancestor of all sampled viruses. This occurrence is different to previous viruses and epidemics and provides some advantages for the development of a rational and scalable classification scheme. Specifically, setting the “reference sequence” to be the “root sequence” forms a natural starting point, as direct comparisons in the number and position of mutations can be made with respect to the root sequence.

During the early phase of the pandemic, it will be possible to unambiguously assign a genome to a lineage through the presence/absence of particular sets of mutations. However, a central component of a useful nomenclature system is that it focuses on those virus lineages that contribute most to global transmission and genetic diversity. Hence, rather than naming every new possible lineage, classification should focus on those that have exhibited onward spread in the population, particularly those that have seeded an epidemic in a new location. For example, the large epidemic in Lombardy, northern Italy, thought to have begun in early February (Zehender et al. 2020), has since been disseminated to other locations in northern Europe and elsewhere.

Further, because SARS-CoV-2 genomes are being generated continuously and at a similar pace to changes in virus transmission and epidemic control efforts, we expect to see a continual process of lineage generation and extinction through time. Rather than maintaining a cumulative list of all lineages that have existed since the start of the pandemic, it is more prudent to mark lineages as ‘active’, ‘unobserved’, or ‘inactive’, a designation reflecting our current understanding of whether they are actively transmitting in the population or not. Although this will allow us to track those lineages that are contributing most to the epidemic, and so reduce the number of names in use, it is important to keep open the possibility that new lineages will appear through the generation of virus genomes from unrepresented locations or from cases with travel history from such locations. For example, the epidemic in Iran, designated B.4 in our system, was identified via returning travellers to other countries (Eden et al. 2020). Further, lineages that have not been seen for some time may re-emerge after a period of cryptic transmission in a region, as proposed for the epidemic in Washington State, US (Bedford et al. 2020). We choose the term lineages (rather than ‘clades’, ‘genotypes’ or other designations) for SARS-CoV-2 as it captures that they are dynamic, rather than relying on a static and exclusive hierarchical structure.

We propose that major lineage labels begin with a letter. At the root of the phylogeny of SARS-CoV-2 are two lineages that we simply denote as lineages A and B. The earliest lineage A viruses, such as Wuhan/WH04/2020 (EPI_ISL_406801), sampled on 2020-01-05, share two nucleotides (positions 8782 in ORF1ab and 28144 in ORF8) with the closest known bat virus (RaTG13). Different nucleotides are present at those sites in viruses assigned to lineage B, of which Wuhan-Hu-1 (Genbank accession MN908947) sampled on 2019-12-26 is an early representative. Hence, although viruses from lineage B were sequenced and published first (Wu et al. 2020; Zhu et al. 2020; Lu et al. 2020), it is likely (based on current data) that lineage A viruses form the root of the SARS-CoV-2 pandemic phylogeny. At the time of writing, viruses from both lineages A and B are still circulating in many countries around the world, reflecting the exportation of viruses from Hubei to other regions of China and elsewhere before the strict travel restrictions and quarantine measures were imposed there.

To add further lineage designations we downloaded 2685 complete SARS-CoV-2 genomes from the GISAID database (Shu and McCauley 2017) on 1st of April, 2020 and estimated a maximum likelihood tree for these data (see Methods) (Figure 1). Following the criteria outlined above, the tree root was placed on the branch between lineages A and B. We then defined further SARS-CoV-2 lineages, each of which descends from either lineage A or B and is assigned a numerical value (e.g. lineage A.1, or lineage B.2). Lineage designations were made using the following set of conditions:

I. Each descendent lineage should show *phylogenetic evidence* of emergence from an ancestral lineage into another geographically distinct population, implying substantial onward transmission in that population. In the case of a rapidly expanding global lineage the recipient “population” may comprise multiple countries. In the case of large and populous countries it may represent a new region or province. To show *phylogenetic evidence* a new lineage must meet *all* of the following criteria: (a) it exhibits one or more shared nucleotide differences from the ancestral lineage, (b) it comprises at least 5 genomes with >95% of the genome sequenced, (c) genomes within the lineage exhibit at least one shared nucleotide change among them, and (d) a bootstrap value >70% for the lineage defining node. Importantly, criterion (c) helps to focus attention only on lineages with evidence of on-going transmission.
II. The lineages identified in step (I) can themselves act as ancestors for virus lineages that then emerge in other geographic areas or at later times, provided they satisfy criteria a-d above. This results in a new lineage designation (e.g. *A.1.1*).
III. The iterative procedure in step II can proceed for a maximum of 3 sublevels (e.g. A.1.1.1) after which new descendent lineages are given a letter (in English alphabetical sequence from C - so A.1.1.1.1 would become C.1 and A.1.1.1.2 would become *C.2*. The rationale for this is that the system is intended only for tracking currently circulating lineages, hence we do not try to capture the entire history of a lineage in its label (that complete history can be obtained by reference to a phylogeny).
IV. All sequences are assigned to one lineage. For example, if a genome does not meet the criteria for inclusion in a “higher level” lineage (e.g. A.1.2, B.1.3.5) then it is automatically classified into the lowest level for which it does meet the inclusion criteria, which ultimately is “A” or B”.

**Figure 1.**
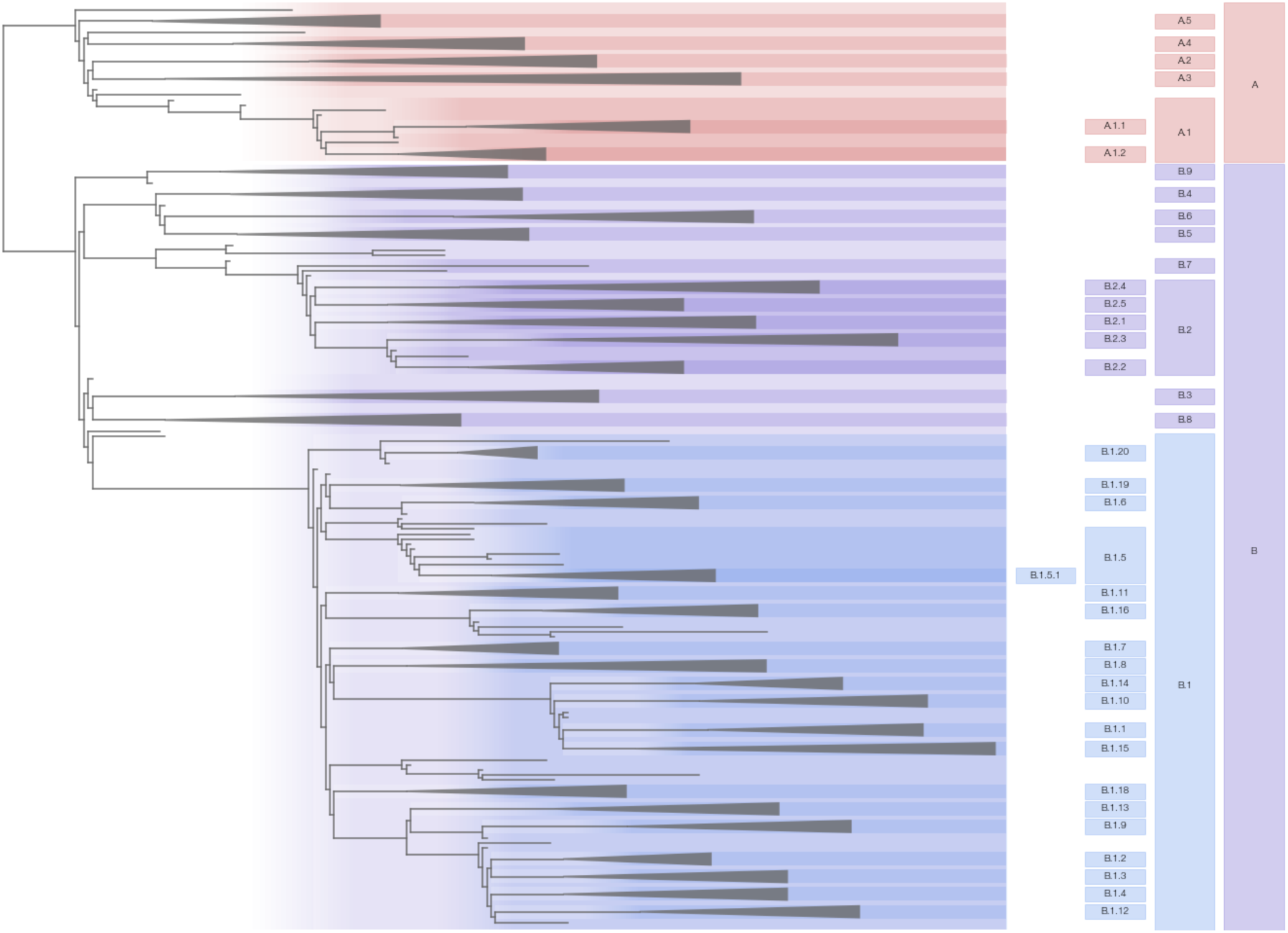
Maximum likelihood phylogeny of globally sampled sequences of SARS-CoV-2 downloaded from the GISAID database (http://gisaid.org) on April 9th 2020. Five representative genomes are included from each of the defined lineages. The tree is rooted on the branch separating lineages A and B sequences as described in the text. The largest lineages that are defined by our proposed nomenclature system are highlighted with coloured areas and labelled on the right. The remaining lineages defined by the nomenclature system are denoted by triangles. The scale bar represents approximately one nucleotide change within the coding region of the genome.

Using this scheme we identified 5 lineages derived from lineage A (denoted lineage A.1 – A.5) and 8 lineages derived from lineage B (B.1 – B.8). We also identified additional descendant lineages, 2 from lineage A.1 (A.1.1 and A.1.2), 20 for lineage B.1 (B.1.1 to B.1.20), and 5 for B.2 (B.2.1 to B.2.5). We are not yet able to further subdivide the other lineages, even though some of these contain very large numbers of genomes. This is because many parts of the world experienced numerous imported cases followed by exponential growth in local transmission. We provide descriptions of these initial lineages, including their geographical locations and time span of sampling, in Table 1. We have also tried to be flexible with the criteria where, for example, the bootstrap value is below 70% but there is strong prior evidence that the lineage exists and is epidemiologically important. In particular, the Italian epidemic comprises two large lineages in our scheme – B.1 and B.2 – reflecting genomes from Italy as well as from large numbers of travellers from these regions and that fall into both lineages.

**Table 1.** Proposed lineage nomenclature for SARS-CoV-2. See https://github.com/hCoV-2019/lineages/ for full details of the sequences assigned to each lineage.

| Lineage | Bootstrap | Countries | Date range | Comments |
| --- | --- | --- | --- | --- |
| A |  |  |  | Hubei, China and travellers from China. |
| A.1 | 100% | Australia, Canada, Iceland, USA | Feb-20, Mar-25 | Primary outbreak in Washington State, USA |
| A.1.1 | 100% | Iceland | Mar-11, Mar-18 |  |
| A.1.2 | 94% | Australia, Canada, USA | Mar-07, Mar-21 |  |
| A.2 | 100% | 12 countries | Feb-25, Mar-26 | Predominantly Spanish lineage |
| A.3 | 100% | Australia, China, New Zealand, Hong Kong, USA, Vietnam | Jan-15, Mar-25 |  |
| A.4 | 100% | Netherlands, USA | Mar-08, Mar-26 |  |
| A.5 | 100% | Australia, Chile, Colombia, Portugal, Spain, USA | Feb-23, Mar-23 |  |
| B |  |  |  | Hubei, China and travellers from China. |
| B.1 | 13% | 43 countries | Feb-20, Apr-01 | With B.2, comprises the large Italian outbreak |
| B.1.1 | 100% | Congo DR, Denmark, Finland, Iceland, Russia, UK | Mar-02, Mar-30 |  |
| B.1.2 | 100% | Canada, USA | Mar-11, Mar-29 |  |
| B.1.3 | 100% | Canada, China, USA | Mar-12, Mar-23 |  |
| B.1.4 | 36% | Algeria, France, Portugal | Mar-02, Mar-19 |  |
| B.1.5 | 100% | 18 countries | Feb-27, Mar-29 |  |
| B.1.6 | 83% | Australia, Belgium, Congo DR | Mar-04, Mar-17 |  |
| B.1.7 | 100% | Australia, Iceland, UK | Feb-22, Mar-29 |  |
| B.1.8 | 100% | Germany, Iceland, Netherlands, Taiwan | Mar-04, Mar-19 |  |
| B.1.9 | 100% | Belgium, Congo DR | Mar-14, Mar-31 |  |
| B.1.10 | 96% | Denmark, Finland, Luxembourg, Norway, UK | Feb-29, Mar-14 |  |
| B.1.11 | 100% | Iceland, UK, USA | Mar-07, Mar-22 |  |
| B.2 | 42% | 20 countries | Jan-25, Mar-25 | With B.1, comprises the large Italian outbreak |
| B.2.1 | 100% | 11 countries | Feb-09, Mar-25 |  |
| B.2.2 | 100% | Australia, Belgium, Iceland, UK, USA | Feb-14, Mar-25 |  |
| B.3 | 100% | 10 countries | Feb-25, Mar-13 |  |
| B.3.1 | 100% | UK | Mar-13, Mar-18 |  |
| B.4 | 100% | 11 countries | Jan-18, Mar-14 | Numerous travelers from Iran |
| B.4.1 | 100% | Australia, Canada, Georgia, Kuwait, Norway, UK | Feb-26, Mar-06 |  |
| B.4.2 | 61% | Australia | Feb-29, Mar-11 |  |
| B.5 | 92% | Japan | Feb-10, Feb-20 |  |
| B.6 | 100% | Australia, Canada, Malaysia, Saudi Arabia, USA | Mar-05, Mar-20 |  |
| B.7 | 99% | Canada, Hong Kong | Jan-30, Mar-04 |  |
| B.8 | 81% | Iceland, Portugal, UK | Mar-02, Mar-25 |  |
| B.9 | 100% | Canada, China, USA | Jan-23, Feb-05 |  |

A unique and important aspect of our proposed nomenclature is that the status of the currently circulating lineages be assessed at regular intervals, with decisions made about identifying new lineages and flagging those we believe are likely be “unobserved” or “inactive” because none of their members have been sequenced for a considerable time (>1 month for “unobserved” and >3 months for “inactive”). When visualising the epidemic we suggest that these lineages should be no longer labelled to reduce both the number of names in circulation and visual noise, and to focus on the current epidemiological situation.

While we regard this proposed nomenclature as practical and robust, it is important to recognise that phylogenetic inference carries statistical uncertainty and much of the available genome data is noisy, with incomplete genome coverage and errors arising from the amplification and sequencing processes. We have proposed a genome coverage threshold for proposing new lineages (see above), and we further suggest that sequences are not ascribed a lineage designation unless the genome coverage of that sequence exceeds 70% of the coding region. As noted above, when SARS-CoV-2 genetic diversity is low during the early pandemic period, there will be a direct association between lineage assignation and the presence of particular sets of mutations (with respect to the root sequence). This should help with the development of algorithmic genome labelling approaches. This task will become more complex, but still tractable, as SARS-CoV-2 genetic diversity accumulates, increasing the chance of both homoplasies and reverse mutations. Classification algorithms based on analysing patterns of mutations may be practical if they are frequently cross-checked and validated against phylogenetic estimations.

Coronaviruses also frequently recombine, meaning that a single phylogenetic tree may not always adequately capture the evolutionary history of the SARS-CoV-2. Therefore, the assignment of some viruses may be ambiguous and may later need to be reassigned. We believe this will not detract from the utility of the system, and as it is intended as a tool for real-time genomic epidemiology, these re-assignment events will quickly become unimportant. In addition, if recombinant lineages arise, exhibit onward spread, and satisfy the requirements for lineage designation outlined above, then they will be assigned the next available alphabetical prefix irrespective of what their parent lineages are.

Finally, while we believe that our proposed lineage nomenclature will greatly assist those working with COVID-19, we do not see it as exclusive to other naming systems, particularly those that are specifically intended to track lineages circulating within individual countries for which a finer scale will be helpful. We envisage, however, that the general approach described here may be readily adopted for these purposes, and also for other viral epidemics where real-time genomic epidemiology is being undertaken.

## Methods

We downloaded all SARS-CoV-2 genomes (at least 29,000bp in length) from GISAID on April 9th 2020. We trimmed the 5’ and 3’ untranslated regions and retained those genomes with at least 95% coverage of the reference genome (Wuhan-Hu-2019, GenBank accession MN908947). We aligned these sequences using MAFFT’s FFT-NS-2 algorithm and default parameter settings (Katoh et al. 2002). We then estimated a maximum likelihood tree using IQ-TREE 2 (Minh et al. 2020) using the GTR+Γ model of nucleotide substitution (Tavaré 1986; Yang 1994), default heuristic search options, and ultrafast bootstrapping with 1000 replicates (Minh, Nguyen, and von Haeseler 2013).

The maximum likelihood tree annotated with the lineage designations, along with a table providing the lineage designation for each genome in the data set, is available for download at https://github.com/hCoV-2019/lineages/. We also provide a high-resolution PDF figure of the entire tree labelled with lineages. These will be updated on a regular basis.

## Notes

https://github.com/hCoV-2019/pangolin

### Competing Interest Statement

The authors have declared no competing interest.

