## Supplementary Table 1 for "A dynamic nomenclature proposal for SARS-CoV-2 to assist genomic epidemiology"

### Supplementary Material

PDF version of tree:

https://www.dropbox.com/s/h09n1to7v9rrwzv/global_lineages_tree.pdf?dl=0

| lineage | boostrap | countries | date range | number of taxa |
|---|---|---|---|---|
| **A** | 69 | Congo; Senegal; South_Korea; USA; Spain; China; Belgium; Australia; Germany; India; UK; Malaysia; Japan; Singapore; Taiwan | Mar-22 Jan-05 | 124 |
| **A.1** | 100 | Virginia; USA; Iceland; Australia; Canada | Mar-25 Feb-20 | 501 |
| **A.1.1** | 100 | Iceland | Mar-18 Mar-11 | 11 |
| **A.1.2** | 91 | USA; Canada; Australia | Mar-21 Mar-07 | 13 |
| **A.2** | 100 | Senegal; USA; Portugal; Spain; Greece; France; Australia; Brazil; UK; Chile; Georgia; Netherlands | Mar-26 Feb-25 | 60 |
| **A.3** | 95 | USA; China; Australia; Vietnam; Hong_Kong; New_Zealand | Mar-25 Jan-15 | 48 |
| **A.4** | 100 | USA; Netherlands | Mar-26 Mar-08 | 18 |
| **A.5** | 100 | USA; Portugal; Australia; Chile; Spain; Colombia | Mar-23 Feb-23 | 15 |
| **B** | 69 | Sweden; Pakistan; Vietnam; Japan; Cambodia; Portugal; Ireland; Iceland; Thailand; France; India; Nepal; Singapore; Spain; Canada; New_Zealand; USA; Belgium; Australia; Germany; Malaysia; Ecuador; Finland; Georgia; Hong_Kong; Taiwan; South_Korea; Slovakia; China; Norway; UK; Italy; Netherlands | Mar-28 Dec-24 | 535 |
| **B.1** | 47 | Greece; Austria; Russia; Nigeria; Japan; Vietnam; Denmark; Mexico; Portugal; Ireland; Iceland; France; India; Switzerland; Brazil; Chile; Senegal; Colombia; Slovenia; Spain; Canada; South_Africa; New_Zealand; Congo; USA; Saudi_Arabia; Czech_Republic; Belgium; Australia; Germany; Finland; Georgia; Argentina; Taiwan; Slovakia; Norway; UK; Hungary; Luxembourg; Italy; Peru; Netherlands | Apr-01 Feb-20 | 1970 |
| **B.1.1** | 93 | Congo; Iceland; Russia; UK; Finland; Denmark | Mar-30 Mar-02 | 64 |
| **B.1.2** | 54 | USA; Canada | Mar-29 Mar-11 | 27 |

| | | | | |
|---|---|---|---|---|
| **B.1.3** | 90 | USA; Canada; China | Mar-23<br>Mar-12 | 49 |
| **B.1.4** | 84 | France; Portugal; Algeria | Mar-19<br>Mar-02 | 9 |
| **B.1.5** | 16 | USA; Portugal; Spain; Iceland; France; Australia; Austria; Switzerland; Norway; UK; Hungary; Luxembourg; Finland; Georgia; Italy; Panama | Mar-29<br>Feb-27 | 104 |
| **B.1.5.1** | 100 | Iceland | Mar-18<br>Mar-12 | 47 |
| **B.1.6** | 80 | Belgium; Australia; Congo | Mar-31<br>Mar-11 | 13 |
| **B.1.7** | 100 | UK; Australia; Iceland | Mar-29<br>Feb-22 | 36 |
| **B.1.8** | 100 | Taiwan; Netherlands; Germany; Iceland | Mar-19<br>Mar-04 | 17 |
| **B.1.9** | 100 | Congo; Belgium | Mar-31<br>Mar-14 | 12 |
| **B.1.10** | 100 | UK; Canada; South_Africa; Iceland | Apr-01<br>Mar-06 | 12 |
| **B.1.11** | 23 | UK; USA; Australia; Iceland | Mar-30<br>Mar-07 | 20 |
| **B.1.12** | 88 | Luxembourg; Belgium | Mar-31<br>Mar-04 | 19 |
| **B.1.13** | 78 | UK; Australia | Mar-28<br>Mar-09 | 16 |
| **B.1.14** | 100 | Iceland | Mar-18<br>Mar-06 | 11 |
| **B.1.15** | 13 | UK; Belgium | Mar-25<br>Mar-01 | 12 |
| **B.1.16** | 75 | Belgium | Mar-20<br>Mar-17 | 5 |
| **B.1.18** | 34 | Belgium; Netherlands | Mar-25<br>Mar-09 | 12 |
| **B.1.19** | 67 | Luxembourg; USA | Mar-25<br>Mar-18 | 11 |
| **B.1.20** | 90 | UK | Mar-30<br>Feb-26 | 9 |
| **B.2** | 13 | Taiwan; USA; Portugal; Ireland; Iceland; Spain; France; Belgium; Australia; Austria; Brazil; UK; China; Korea; Luxembourg; Norway; Finland; Switzerland; Singapore; Netherlands | Mar-29<br>Feb-25 | 167 |

| | | | | |
|---|---|---|---|---|
| **B.2.1** | 13 | Congo; USA; Portugal; Ireland; Iceland; Belgium; Australia; Germany; Brazil; UK; Finland; Canada; Netherlands | Mar-29<br>Feb-09 | 180 |
| **B.2.2** | 100 | USA; Iceland; Belgium; Australia; UK | Mar-28<br>Feb-25 | 31 |
| **B.2.3** | 27 | Belgium | Mar-31<br>Mar-07 | 6 |
| **B.2.4** | 97 | UK; Australia; New_Zealand | Mar-28<br>Mar-11 | 12 |
| **B.2.5** | 100 | Slovenia; UK; Australia; Spain | Mar-27<br>Feb-27 | 11 |
| **B.3** | 97 | USA; Portugal; Iceland; Belgium; Australia; Austria; Brazil; UK; Germany; Finland; Senegal; Poland; Canada; Netherlands | Mar-31<br>Feb-25 | 204 |
| **B.4** | 100 | Congo; Taiwan; USA; Kuwait; New_Zealand; China; Australia; Austria; Germany; Norway; UK; Pakistan; Georgia; Canada; Netherlands | Mar-30<br>Jan-18 | 90 |
| **B.5** | 86 | Japan; Australia | Feb-24<br>Feb-10 | 18 |
| **B.6** | 100 | USA; Saudi_Arabia; Australia; Malaysia; Canada; South_Africa | Mar-31<br>Mar-05 | 22 |
| **B.7** | 13 | Canada; Hong_Kong | Mar-04<br>Jan-30 | 40 |
| **B.8** | 70 | UK; Portugal; Iceland | Mar-29<br>Mar-03 | 34 |
| **B.9** | 20 | Netherlands | Mar-09<br>Mar-08 | 6 |
